# Physicochemical compatibility and stability of urapidil-propofol admixtures during simulated Y-site Administration

**DOI:** 10.64898/2026.09.07.749885

**Authors:** Yantao Yue, Liguo Wang, Yuanpei Li, Jingduo Zhao, Lijie Lou, Shijie Feng, Liang Zhao

## Abstract

**Background/Objectives:** Urapidil with propofol is clinically efficient against elevated blood pressure during sedation. However, their physicochemical compatibility and emulsion stability upon continuous infusion remain unclear. This study aimed to evaluate different mixing ratios and diluents, thereby proving the safety limits for their co-administration.

**Methods:** Urapidil solutions prepared with either sodium chloride (NS) or glucose injection (GS), and emulsified with propofol at different ratios (v/v), were stored for 12 h. Physical compatibility was assessed by visual inspection, pH, osmolality, mean droplet diameter (MDD), polydispersity index (PDI), zeta potential, and percentage of fat globules larger than 5 μm (PFAT_5_). Chemical stability was quantified using high-performance liquid chromatography.

**Results:** pH and osmolality stabilized. Urapidil hydrochloride and propofol contents remained pure at > 95%, MDD was < 500 nm, and PDI was < 0.2. Urapidil proportion in NS was significantly negatively correlated with the zeta potential. PFAT_5_ was > 0.05% after 2‒8 h. In contrast, in GS at a 1:2 ratio, PFAT_5_ remained < 0.05%, which increased slightly in the 1:1 group at 8 h. PFAT_5_ stabilized in the high-propofol group (10:1) under all conditions.

**Conclusions:** The chemical compatibility of the admixture was acceptable after 12 h of storage. However, physical compatibility was influenced by the mixing ratio, diluent type, and storage time. For clinical Y-site co-administration, a 10:1 mixing ratio or dilution in 5% GS is recommended. Enhanced proportions must be mixed with NS, while continuous infusion time must be < 2 h to mitigate fat embolism risk.

## Introduction

Hypertensive intracerebral hemorrhage (HICH) is a common neurocritical condition associated with extremely high mortality and disability [1]. Marked blood pressure (BP) fluctuations contribute to hematoma expansion, rebleeding, and poor prognosis [2]. Therefore, postoperative management focuses on blood pressure control, intracranial pressure reduction, and cerebral edema alleviation [3]. Urapidil, a selective α1-adrenergic receptor antagonist, reduces peripheral vascular resistance without inducing reflex tachycardia and is a common first-line agent for acute BP management in HICH [4]. To attenuate stress responses and agitation associated with surgical trauma and endotracheal intubation, propofol is a widely used postoperative sedative because of its rapid onset, short recovery time, and ease of titration [5]. Clinical studies in postoperative patients with HICH have shown that combined urapidil and propofol therapy better stabilizes BP and heart rate, reduces intracranial pressure, and improves clinical outcomes compared with urapidil monotherapy [6,7].

However, in clinical practice, critically ill patients often require the simultaneous infusion of multiple medications through a single central venous catheter via Y-site connectors or multiway stopcocks, increasing the risk of physicochemical incompatibility [8]. Such incongruence may cause drug precipitation, discoloration, degradation, or the formation of toxic compounds and insoluble particles, potentially leading to phlebitis, pulmonary embolism, or systemic inflammatory response syndrome [9,10]. Although the pharmacodynamic impacts of combined urapidil-propofol therapy have been reported, data on their physicochemical compatibility and stability during Y-site infusion remain limited. As a lipid emulsion, propofol stability is susceptible to pH, electrolyte concentration, and emulsion destabilization or droplet aggregation induced by co-administered drugs, which may enhance embolism risk [11,12]. In particular, the effects of concomitant drug administration on the percentage of fat globules larger than 5 μm (PFAT_5_), a key indicator of lipid emulsion stability, remain unclear [13,14].

Therefore, based on clinically relevant doses, this study used 0.9% sodium chloride injection (normal saline, NS) and 5% glucose injection (GS) as diluents to prepare propofol‒urapidil admixtures of various ratios. Under simulated conditions of clinical Y-site infusion, the physicochemical compatibility and emulsion stability of the admixtures were systematically evaluated at room temperature over a predefined period. This study provides experimental evidence to support the safe and rational co-administration of these medications in critically ill patients.

## Materials and methods

### Drugs and reagents

Propofol emulsion (20 mL: 0.2 g) was obtained from Cisen Pharmaceutical Co., Ltd. (Jining, China). Urapidil injection (5 mL: 25 mg) was purchased from Fuhe Pharmaceutical Group Co., Ltd. (Heilongjiang, China). 0.9% NS (50 mL) and 5% GS (50 mL) were supplied by Sichuan Kelun Pharmaceutical Co., Ltd. (Chengdu, China). Urapidil hydrochloride, urapidil impurity I (1,3-dimethyl-4-(γ-chloropropylamino)uracil), propofol, propofol impurity I (3,3’,5,5’-tetraisopropyldiphenol-4,4’-diol), and propofol impurity II(2,6-diisopropyl-1,4-benzoquinone) reference standards were bought from the National Institute for Food and Drug Control (Beijing, China). Methanol and acetonitrile of high-performance liquid chromatography (HPLC) grade were used, and all other chemicals were of analytical grade; they were purchased from Shanghai Aladdin Biochemical Technology Co., Ltd. (Shanghai, China).

### Sample preparation

Preparation of urapidil solutions: Four ampoules of urapidil injection were separately diluted with 30 mL of 0.9% NS and 5% GS. After thorough mixing, the solutions were transferred into 50 mL syringes to prepare urapidil-NS and urapidil-GS, each at a final concentration of 2 mg/mL.

Preparation of admixture solutions: Sample preparation was designed to simulate clinical infusion protocols. Based on common infusion rates, propofol was administered at 0.3–4 mg/kg·h (2.1–28 mL/h), whereas urapidil was administered at 5–60 mg/h (2.5–30 mL/h). Accordingly, the potential mixing ratios in the Y-site pathway were determined, and three representative volume ratios of the propofol original solution to the urapidil solution (10:1, 1:1, and 1:10) were selected to reflect varied clinical scenarios. For each ratio and diluent, three independent parallel samples were prepared. Samples were stored at room temperature—25 ± 2 °C—under indoor light exposure. Aliquots were collected at 0, 1, 2, 4, 8, and 12 h, and each parameter was analyzed in triplicate at each time point. Related information is detailed in Table 1.

**Table 1.** Composition of admixture solutions.

| Grou<br>p | Composition | Mixing<br>Ratio(v/v) |
| --- | --- | --- |
| 1 | Propofol original solution + urapidil solution<br>in 0.9% sodium chloride | 10:1 |
| 2 | Propofol original solution + urapidil solution<br>in 0.9% sodium chloride | 1:1 |
| 3 | Propofol original solution + urapidil solution<br>in 0.9% sodium chloride | 1:10 |
| 4 | Propofol original solution + urapidil solution<br>in 5% glucose | 10:1 |
| 5 | Propofol original solution + urapidil solution<br>in 5% glucose | 1:1 |
| 6 | Propofol original solution + urapidil solution<br>in 5% glucose | 1:10 |

### Visual inspection

According to the relevant provisions of the Chinese Pharmacopoeia [15], all admixtures were visually inspected for particles and color changes. Observations were made against a black-and-white contrast background by two independent observers. The admixtures were considered visually compatible if no precipitation, phase separation, or pronounced color changes were observed.

### pH and osmolality measurements

At each predetermined time point, 5 mL was aliquoted for pH measurement at room temperature with an FE28 pH meter (Mettler-Toledo GmbH, Switzerland). Additionally, 60 μL was aliquoted to determine osmolality at room temperature using an SMC 30C-1 osmometer (Tianjin Tianhe Analytical Instrument Co., Ltd., China). Samples were considered stable and physically compatible if the pH change was ≤ 0.2 and the osmolality change was <5% compared with the baseline values, immediately after mixing [16].

### Droplet size and zeta potential measurements

At each time point, samples were collected and diluted 150-fold with purified water filtered through a 0.22 μm membrane based on the propofol concentration. The mean droplet diameter (MDD) and zeta potential (ζ) of the emulsions were measured with a Zetasizer Lab ZS (Malvern Instruments, Malvern, UK) via dynamic light scattering and electrophoretic light scattering, respectively. Droplet size distribution was evaluated employing the polydispersity index (PDI). According to the requirements of the Chinese Pharmacopoeia and the United States Pharmacopoeia (USP) for intravenous lipid emulsions, the intensity-weighted MDD of lipid droplets in an admixture should not exceed 500 nm [17].

### Percentage of fat globules larger than 5 µm (PFAT_5_) determination

At each time point, samples were collected and analyzed using an Accusizer 780 APS particle sizer (PSS, Santa Barbara, CA, USA). The number and size of oil droplets > 1 µm were measured via the light obscuration/single-particle optical sensing (LO/SPOS) technique. PFAT_5_ was calculated as the weighted volume of droplets > 5 µm relative to the volume of the total dispersed phase. Measurements were in extinction (EXT) mode with a detection range of 1.8–50 µm. The dilution flow rate was set at 60 mL/min, with a target particle concentration of 2000 particles/mL and a sample injection volume of 0.98 mL. Droplet concentration and size distribution were recorded to calculate the PFAT_5_. According to the Chinese Pharmacopoeia and USP, lipid emulsions are considered physically stable when PFAT_5_ values are < 0.05%.

### Chromatographic conditions

Chromatography was performed on an LC-15C HPLC System (Shimadzu Corporation, Japan) equipped with an Agilent InfinityLab Poroshell 120 EC-C18 column (4.6 mm × 150 mm, 2.7 µm). The mobile phase consisted of acetonitrile (A) and sodium dihydrogen phosphate solution (B). The latter was prepared by dissolving 2.76 g of sodium dihydrogen phosphate monohydrate in 900 mL of water, adjusting the pH to 3.0 with 85% phosphoric acid, and diluting to 1,000 mL with water. Gradient elution was performed: 0–7 min, 14% A; 7–8 min, 14%‒40% A; 8–20 min, 40% A; 20–35 min, 40%‒70% A; 35–36 min, 70%‒90% A; 36–40 min, 90% A; 40–41 min, 90%‒14% A. The flow rate was 1.0 mL/min, and the column temperature was 38℃. The detection wavelength was 268 nm, and the injection volume was 10 µL.

### Methodological validation

#### Sample preparation

An aliquot (0.4 mL) of each admixture was accurately transferred into a 5 mL centrifuge tube, and 2.4 mL of methanol was added. The mixture was vortexed for 30 s and centrifuged at 233.3 s⁻¹ (14,000 rpm) for 10 min (r = 6 cm) with a high-speed centrifuge. Subsequently, 2 mL of the supernatant was transferred into a 5 mL volumetric flask and diluted to the required volume with methanol. The solution obtained was filtered through a 0.22 μm microporous membrane before analysis.

#### Specificity

Chromatograms of the sample, reference standard, and blank vehicle solutions were obtained under the conditions described in Section 2.7. The corresponding peaks in the samples and references showed identical retention times, and propofol, urapidil hydrochloride, and their related impurities were well separated. The blank vehicle did not exhibit any interfering peaks at the corresponding retention times, demonstrating the specificity of the analytical method. The chromatograms are presented in Fig 1.

**Fig 1.**
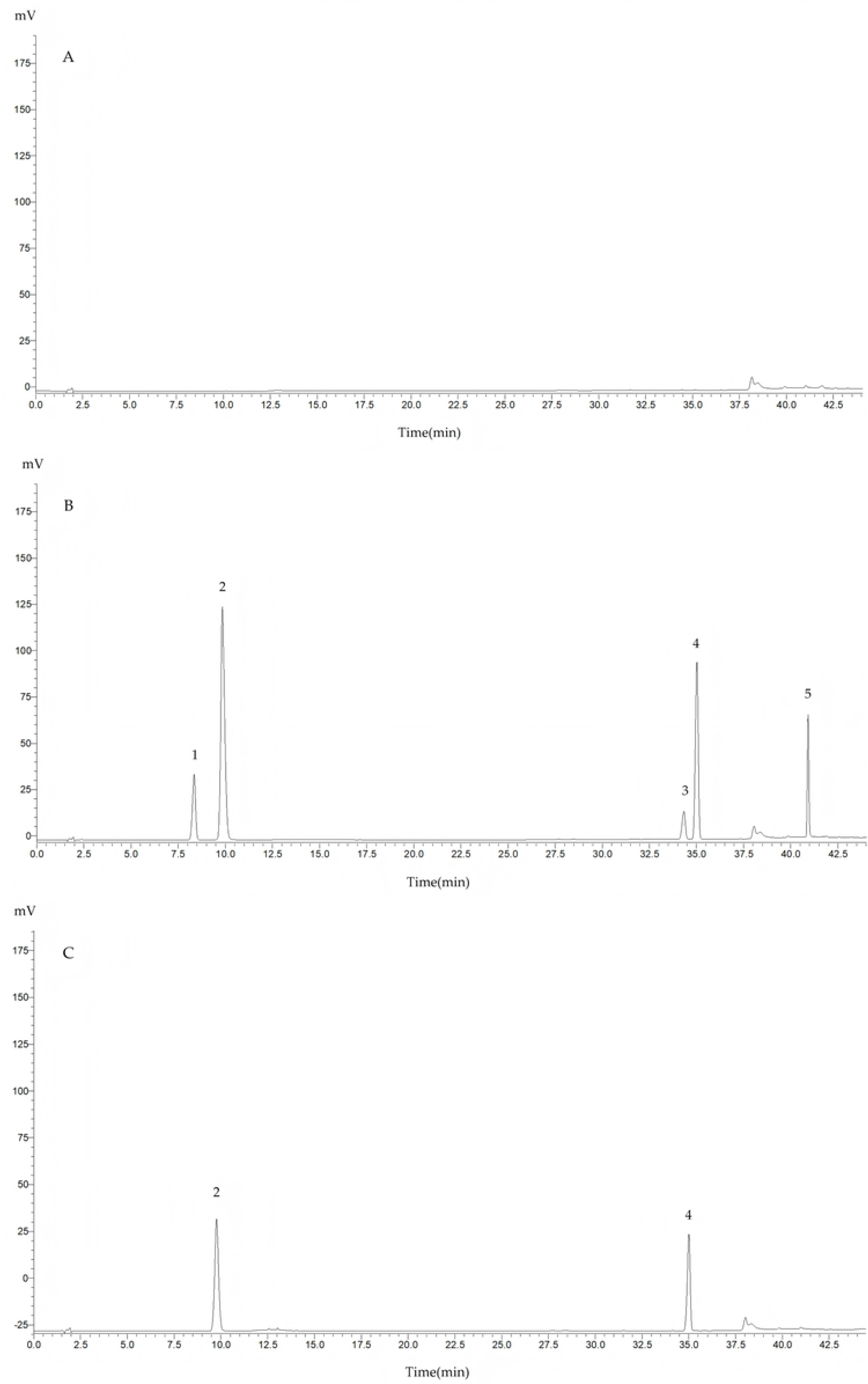
High-performance liquid chromatograms. (A) blank control; (B) mixed reference standard solution; (C) test sample solution (1. urapidil impurity I; 2. urapidil hydrochloride; 3. propofol impurity II; 4. propofol; 5. propofol impurity I).

#### Calibration curves

Aliquots of the five reference standards were accurately transferred and diluted with methanol to prepare standard working solutions at six concentrations: Propofol, 2.09‒209.0 μg/mL; propofol impurity I 0.45‒45.0 μg/mL; propofol impurity II 0.5‒50.0 μg/mL; urapidil hydrochloride, 2.65‒106.0 μg/mL; and urapidil impurity I, 0.375‒15.0 μg/mL. These were chromatographically analyzed under the conditions described in Section 2.7, and peak areas were recorded. Calibration curves were constructed by plotting peak area (Y) vs. concentration (X). The corresponding regression equations and linear ranges were, Propofol, Y1 = 4,761.5X1 − 566.39 (r = 0.9999), with a linear range of 2.09–209.0 μg/mL; Propofol impurity I, Y2 = 36,253X2 + 18.311 (r = 0.9999), with a linear range of 0.45–45.0 μg/mL; Propofol impurity II, Y3 = 37,553X3 + 217.72 (r = 0.9999), with a linear range of 0.5–50.0 μg/mL; Urapidil hydrochloride, Y4 = 33,363X4 + 37,581 (r = 0.9998), with a linear range of 2.65–106.0 μg/mL; Urapidil impurity I, Y5 = 28,654X5 − 1,320.7 (r = 0.9999), with a linear range of 0.375–15.0 μg/mL.

#### Precision

Admixtures were injected six consecutive times; the peak areas of each component were recorded, and the relative standard deviations (RSDs) were calculated. The RSDs of propofol and urapidil hydrochloride were 0.31% and 0.47% (n = 6), respectively, suggesting good instrumental precision.

#### Repeatability

Six parallel samples of the admixture were prepared per the method described for Group 2 in Section 2.2. After preparation as described in Section 2.8.1, the samples were analyzed, and the peak areas of propofol and urapidil hydrochloride were recorded to calculate the RSDs. The RSDs of the propofol and urapidil hydrochloride peak areas were 1.96% and 1.93% (n = 6), respectively, indicating satisfactory method repeatability.

#### Sample recovery test

Nine aliquots of the admixture containing known propofol and urapidil hydrochloride concentrations were prepared. Reference standard solutions at low, medium, and high levels were accurately added, with three replicates for each. After processing, the samples were analyzed to determine the recoveries of propofol and urapidil hydrochloride. The results are presented in Table 2.

**Table 2.** Results of sample recovery test for propofol and urapidil hydrochloride (n=9)

| Component | Sample<br>amount<br>(μg) | Added<br>amount(μg) | Measured<br>amount(μg) | Recovery(%) | Average<br>recovery(%) | RSD(%) |
| --- | --- | --- | --- | --- | --- | --- |
| Urapidil | 3.015 | 2.650 | 5.522 | 97.48 | 98.93 | 1.38 |
|  | 3.015 | 2.650 | 5.679 | 100.20 |  |  |
|  | 3.015 | 2.650 | 5.614 | 99.10 |  |  |
|  | 3.015 | 10.600 | 13.258 | 97.38 | 98.95 | 1.49 |
|  | 3.015 | 10.600 | 13.502 | 99.17 |  |  |
|  | 3.015 | 10.600 | 13.656 | 100.30 |  |  |
|  | 3.015 | 31.800 | 34.264 | 98.42 | 98.47 | 1.86 |
|  | 3.015 | 31.800 | 34.590 | 99.35 |  |  |
|  | 3.015 | 31.800 | 33.997 | 97.65 |  |  |
| Propofol | 5.986 | 10.450 | 16.332 | 99.37 | 99.59 | 0.63 |
|  | 5.986 | 10.450 | 16.287 | 99.10 |  |  |
|  | 5.986 | 10.450 | 16.490 | 100.30 |  |  |
|  | 5.986 | 20.900 | 26.548 | 98.74 | 98.03 | 0.91 |
|  | 5.986 | 20.900 | 26.435 | 98.32 |  |  |
|  | 5.986 | 20.900 | 26.086 | 97.02 |  |  |
|  | 5.986 | 62.700 | 67.965 | 98.95 | 99.37 | 0.64 |
|  | 5.986 | 62.700 | 68.036 | 99.05 |  |  |
|  | 5.986 | 62.700 | 68.754 | 100.10 |  |  |

#### Stability of drug solutions

Urapidil-NS, urapidil-GS, and propofol stock solutions prepared as described in Section 2.2 were transferred into 50 mL syringes and stored at room temperature for 12 h. They were sampled at 0, 1, 2, 4, 8, and 12 h. For urapidil solutions, 50 μL aliquots were accurately transferred, diluted with 1 mL methanol, and 5 μL aliquots were injected for analysis. For propofol stocks, 140 μL were accurately pipetted, mixed with 1,820 μL methanol, and centrifuged at 233.3 s⁻¹ (14,000 rpm) for 10 min. Subsequently, 800 μL of the supernatant was collected, diluted with 400 μL of methanol, and mixed thoroughly before HPLC analyses, under the conditions described in Section 2.7, and the peak areas were recorded. The RSDs of the peak areas for urapidil-NS, urapidil-GS, and RSD propofol were 1.52%, 1.52%, and 1.36%, respectively, indicating stability of the drug contents over the 12-h storage period.

### Determination of drug contents and related compounds

Sample solutions prepared as described in Section 2.2 were chromatographically analyzed under the conditions described in Section 2.7. Peak areas of each component were recorded to calculate the drug contents; those at 0 h were defined as 100%, and the relative contents (%) at subsequent time points were calculated accordingly.

When impurity peaks corresponding to urapidil impurity I, propofol impurities I and II were detected in the chromatograms, according to the relevant requirements of the 2020 edition of the Chinese Pharmacopoeia, the contents of propofol impurities I and II should each not exceed 0.1% of the labeled propofol content, and the peak area of urapidil impurity I must not exceed 0.5% of the reference standard peak area.

### Statistical analysis

All statistical analyses were performed using SPSS version 25.0. Data were presented as mean ± standard deviation. The Shapiro-Wilk test was used to assess normality. For data conforming to a normal distribution, a two-factor repeated-measures analysis of variance was performed. Mauchly’s test of sphericity was applied, and the Greenhouse-Geisser correction was employed when the sphericity assumption was violated. Mixing ratio and diluent type were considered inter-subject factors, while time was considered an intra-subject factor. Main and interaction effects were evaluated, and Bonferroni-adjusted post hoc comparisons were performed when effects were significant; statistical significance was defined as p < 0.05.

## Results

### Visual inspection

Under storage at room temperature (25 ± 2 °C) with 12 h of indoor light exposure, all six groups of propofol-urapidil admixtures remained white, homogeneous emulsions. No visible signs of incompatibility, including phase separation, precipitation, color change, or oiling out, were observed.

### pH and osmolality

The original propofol solution had a pH of 7.52 and an osmolality of 299 mmol/kg. Urapidil-NS had a pH of 5.97 and an osmolality of 777 mmol/kg, whereas urapidil-GS had corresponding values of 6.15 and 763 mmol/kg. The admixture exhibited pH values ranging from 6.11 to 6.74, and the osmolality ranged from 341 to 758 mmol/kg. Following mixing, pH values fell between those of the two original solutions, whereas osmolality varied according to the diluent and mixing ratio. After 12 h of storage, pH changed by less than 0.2 and osmolality by less than 5% in all groups, indicating physicochemical stability throughout the study period (Table 3).

**Table 3.** Results of pH and osmolality for each experimental group (x̅±s)

| Grou<br>p | pH |  |  |  |  |  | Osmolality (mmol/kg) |  |  |  |  |  |
| --- | --- | --- | --- | --- | --- | --- | --- | --- | --- | --- | --- | --- |
|  | 0h | 1h | 2h | 4h | 8h | 12h | 0h | 1h | 2h | 4h | 8h | 12h |
| 1 | 6.60± | 6.61± | 6.62± | 6.61± | 6.61± | 6.61± | 343± | 345± | 345± | 347± | 348± | 353± |
|  | 0.01 | 0.01 | 0.01 | 0.01 | 0.01 | 0.01 | 1 | 3 | 1 | 2 | 2 | 2 |
| 2 | 6.11± | 6.12± | 6.13± | 6.11± | 6.12± | 6.12± | 544± | 544± | 544± | 547± | 551± | 554± |
|  | 0.01 | 0.01 | 0.01 | 0.03 | 0.01 | 0.01 | 1 | 2 | 0 | 2 | 1 | 2 |
| 3 | 6.03± | 6.00± | 5.99± | 5.99± | 5.99± | 6.01± | 723± | 722± | 726± | 728± | 734± | 758± |
|  | 0.04 | 0.02 | 0.01 | 0.01 | 0.01 | 0.01 | 1 | 2 | 2 | 1 | 3 | 1 |
| 4 | 6.72± | 6.74± | 6.72± | 6.71± | 6.72± | 6.71± | 342± | 342± | 341± | 342± | 341± | 344± |
|  | 0.01 | 0.01 | 0.01 | 0.00 | 0.00 | 0.01 | 1 | 1 | 2 | 1 | 2 | 4 |
| 5 | 6.25± | 6.28± | 6.24± | 6.24± | 6.24± | 6.23± | 540± | 539± | 540± | 538± | 537± | 541± |
|  | 0.01 | 0.01 | 0.01 | 0.01 | 0.01 | 0.01 | 2 | 4 | 4 | 5 | 2 | 3 |
| 6 | 6.12± | 6.11± | 6.11± | 6.11± | 6.12± | 6.12± | 723± | 724± | 723± | 723± | 722± | 727± |
|  | 0.02 | 0.02 | 0.02 | 0.02 | 0.01 | 0.01 | 3 | 4 | 3 | 3 | 1 | 2 |

### Droplet size and zeta potential

The MDD did not vary significantly among groups with different mixing ratios using the same diluent or conversely (P > 0.05). Over time, MDD altered significantly only in Group 1 (P < 0.05), but not among the remaining groups. The MDDs of all admixtures ranged from 169.1 to 182.7 nm, well below the 500 nm limit specified by the USP and the Chinese Pharmacopoeia. PDI values ranged from 0.04 to 0.11, with no marked differences between groups. All PDI values were below 0.2, indicating a highly uniform droplet size distribution without significant aggregation or phase separation. Neither storage time nor mixing ratio remarkably affected MDD (Table 4).

**Table 4.**
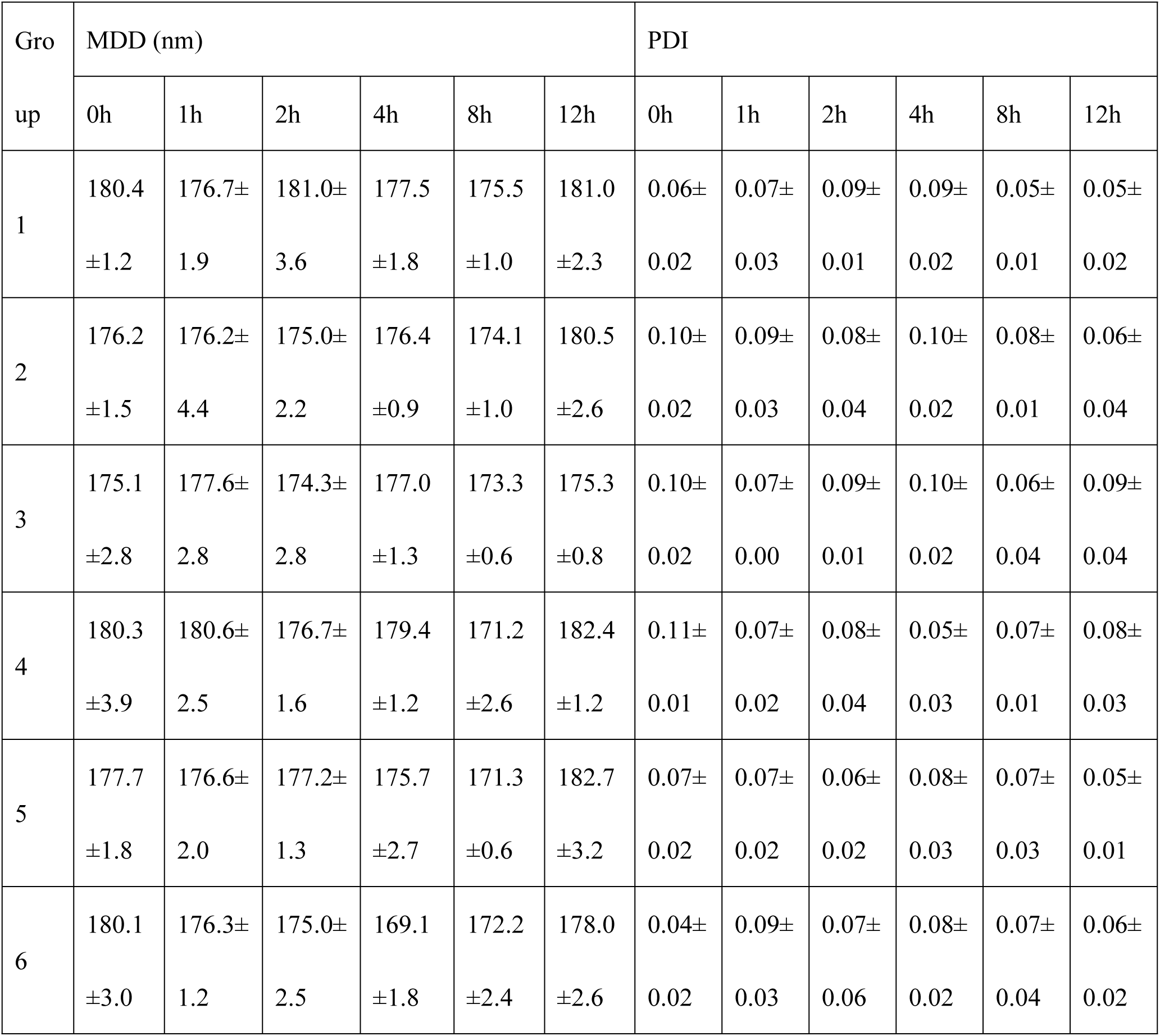
Results of mean droplet diameter (MDD) and polydispersity index (PDI) for each experimental group (x̅±s)

| Gro<br>up | MDD (nm) |  |  |  |  |  | PDI |  |  |  |  |  |
| --- | --- | --- | --- | --- | --- | --- | --- | --- | --- | --- | --- | --- |
|  | 0h | 1h | 2h | 4h | 8h | 12h | 0h | 1h | 2h | 4h | 8h | 12h |
| 1 | 180.4 | 176.7± | 181.0± | 177.5 | 175.5 | 181.0 | 0.06± | 0.07± | 0.09± | 0.09± | 0.05± | 0.05± |
|  | ±1.2 | 1.9 | 3.6 | ±1.8 | ±1.0 | ±2.3 | 0.02 | 0.03 | 0.01 | 0.02 | 0.01 | 0.02 |
| 2 | 176.2 | 176.2± | 175.0± | 176.4 | 174.1 | 180.5 | 0.10± | 0.09± | 0.08± | 0.10± | 0.08± | 0.06± |
|  | ±1.5 | 4.4 | 2.2 | ±0.9 | ±1.0 | ±2.6 | 0.02 | 0.03 | 0.04 | 0.02 | 0.01 | 0.04 |
| 3 | 175.1 | 177.6± | 174.3± | 177.0 | 173.3 | 175.3 | 0.10± | 0.07± | 0.09± | 0.10± | 0.06± | 0.09± |
|  | ±2.8 | 2.8 | 2.8 | ±1.3 | ±0.6 | ±0.8 | 0.02 | 0.00 | 0.01 | 0.02 | 0.04 | 0.04 |
| 4 | 180.3 | 180.6± | 176.7± | 179.4 | 171.2 | 182.4 | 0.11± | 0.07± | 0.08± | 0.05± | 0.07± | 0.08± |
|  | ±3.9 | 2.5 | 1.6 | ±1.2 | ±2.6 | ±1.2 | 0.01 | 0.02 | 0.04 | 0.03 | 0.01 | 0.03 |
| 5 | 177.7 | 176.6± | 177.2± | 175.7 | 171.3 | 182.7 | 0.07± | 0.07± | 0.06± | 0.08± | 0.07± | 0.05± |
|  | ±1.8 | 2.0 | 1.3 | ±2.7 | ±0.6 | ±3.2 | 0.02 | 0.02 | 0.02 | 0.03 | 0.03 | 0.01 |
| 6 | 180.1 | 176.3± | 175.0± | 169.1 | 172.2 | 178.0 | 0.04± | 0.09± | 0.07± | 0.08± | 0.07± | 0.06± |
|  | ±3.0 | 1.2 | 2.5 | ±1.8 | ±2.4 | ±2.6 | 0.02 | 0.03 | 0.06 | 0.02 | 0.04 | 0.02 |

The zeta potential of the admixture ranged from −7.01 to −51.80 mV. Significant variations were observed among mixing ratios and between diluents (P < 0.001). Furthermore, the absolute zeta potential decreased with increasing proportions of urapidil. Although these slightly fluctuated during storage, no pronounced or consistent temporal trends were observed (Table 5).

**Table 5.**
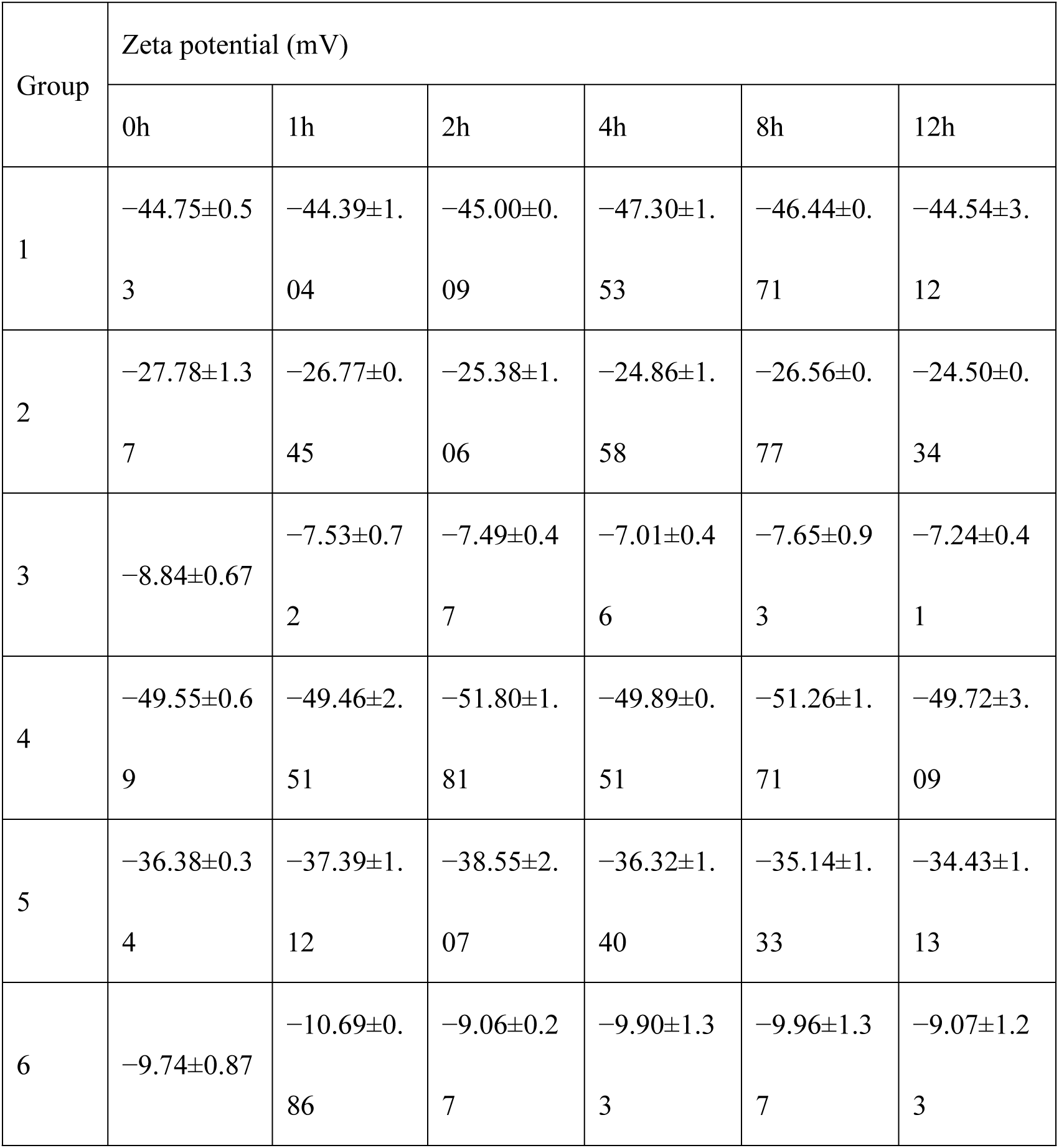
Results of zeta potential for each experimental group (x̅±s)

| Group | Zeta potential (mV) |  |  |  |  |  |
| --- | --- | --- | --- | --- | --- | --- |
|  | 0h | 1h | 2h | 4h | 8h | 12h |
| 1 | -44.75±0.5 | -44.39±1. | -45.00±0. | -47.30±1. | -46.44±0. | -44.54±3. |
|  | 3 | 04 | 09 | 53 | 71 | 12 |
| 2 | -27.78±1.3 | -26.77±0. | -25.38±1. | -24.86±1. | -26.56±0. | -24.50±0. |
|  | 7 | 45 | 06 | 58 | 77 | 34 |
| 3 | -8.84±0.67 | -7.53±0.7 | -7.49±0.4 | -7.01±0.4 | -7.65±0.9 | -7.24±0.4 |
|  |  | 2 | 7 | 6 | 3 | 1 |
| 4 | -49.55±0.6 | -49.46±2. | -51.80±1. | -49.89±0. | -51.26±1. | -49.72±3. |
|  | 9 | 51 | 81 | 51 | 71 | 09 |
| 5 | -36.38±0.3 | -37.39±1. | -38.55±2. | -36.32±1. | -35.14±1. | -34.43±1. |
|  | 4 | 12 | 07 | 40 | 33 | 13 |
| 6 | -9.74±0.87 | -10.69±0. | -9.06±0.2 | -9.90±1.3 | -9.96±1.3 | -9.07±1.2 |
|  |  | 86 | 7 | 3 | 7 | 3 |

### Percentage of PFAT_5_

Storage time, mixing ratio, and diluent type significantly affected PFAT_5_ values (P < 0.001). During the 12-h observation period, PFAT_5_ stabilized in Groups 1, 4, and 6 (P > 0.05), with values consistently below the pharmacopeial safety limit of 0.05%, indicating excellent emulsion stability. In contrast, PFAT_5_ values were enhanced significantly over time in Groups 2, 3, and 5 (P < 0.05), exceeding the safety limit in a temporal manner.

Within the 0.9% NS system, increasing the proportion of urapidil to 1:1 or 1:10 significantly enhanced PFAT_5_ compared with the 10:1 ratio (P < 0.001). Group 2 (1:1, NS) exceeded the pharmacopeial limit at 2 h, whereas Group 3 (1:10, NS) crossed the limit at 8 h. At the same mixing ratio, PFAT_5_ values were markedly higher in the NS than in the GS groups (P < 0.05). Notably, Group 6 (1:10, GS) maintained PFAT_5_ values below the safety limit throughout the 12-h study period, unlike Group 3 (1:10, NS).

These findings indicate that PFAT_5_ exceedance is driven by the combined influence of a high proportion of urapidil, the use of 0.9% NS as the diluent, and prolonged storage. In contrast, elevated proportions of propofol (10:1) or the use of 5% GS preserved emulsion stability over 12 h. Notably, all samples with PFAT_5_ values exceeding the pharmacopeial limit maintained MDD values below 500 nm, indicating that PFAT_5_ was more sensitive and an indicator of earlier emulsion destabilization than MDD (Table 6).

**Table 6.** Results of PFAT_5_ for each experimental group (x̅±s)

| Group | PFAT5 (%) |  |  |  |  |  |
| --- | --- | --- | --- | --- | --- | --- |
|  | 0h | 1h | 2h | 4h | 8h | 12h |
| 1 | 0.007±0.00 | 0.009±0.0 | 0.005±0.0 | 0.009±0.0 | 0.008±0.0 | 0.004±0.0 |
|  | 1 | 02 | 02 | 06 | 03 | 03 |
| 2 | 0.010±0.00 | 0.013±0.0 | 0.169±0.0 | 0.129±0.0 | 0.103±0.0 | 0.070±0.0 |
|  | 3 | 03 | 48 | 35 | 03 | 03 |
| 3 | 0.043±0.02 | 0.026±0.0 | 0.048±0.0 | 0.054±0.0 | 0.103±0.0 | 0.150±0.0 |
|  | 2 | 18 | 14 | 19 | 23 | 09 |
| 4 | 0.005±0.00 | 0.006±0.0 | 0.004±0.0 | 0.006±0.0 | 0.005±0.0 | 0.005±0.0 |
|  | 1 | 01 | 02 | 03 | 02 | 02 |
| 5 | 0.006±0.00 | 0.012±0.0 | 0.016±0.0 | 0.020±0.0 | 0.054±0.0 | 0.043±0.0 |
|  | 0 | 04 | 05 | 02 | 03 | 08 |
| 6 | 0.047±0.01 | 0.044±0.0 | 0.049±0.0 | 0.035±0.0 | 0.039±0.0 | 0.026±0.0 |
|  | 3 | 22 | 18 | 20 | 23 | 14 |

### Drug content and related substances

At room temperature, the relative contents of urapidil hydrochloride and propofol of all admixture groups remained within 95.24%–100.25% and 96.29%–100.05%, respectively, over the 12-h study period. The changes in the contents of both drugs were < 5%, indicating good chemical stability without prominent degradation. Furthermore, impurity peaks corresponding to urapidil impurity I, propofol impurities I or II were not detected in any chromatograms. Except for the main drug and solvent peaks, no additional chromatographic peaks were observed, indicating that none of the degradation products exceeding pharmacopoeial limits were generated during mixing or storage (Table 7).

**Table 7.** Results of drug content determination for each experimental group (x̅±s)

| Gro<br>up | Percentage of initial urapidil concentration (%) |  |  |  |  | Percentage of initial propofol concentration (%) |  |  |  |  |
| --- | --- | --- | --- | --- | --- | --- | --- | --- | --- | --- |
|  | 1h | 2h | 4h | 8h | 12h | 1h | 2h | 4h | 8h | 12h |
| 1 | 99.24± | 98.34± | 98.55± | 96.88± | 96.63± | 99.89± | 100.05 | 99.31± | 98.16± | 97.76± |
|  | 1.07 | 1.82 | 0.74 | 0.65 | 1.05 | 0.49 | ±0.21 | 0.40 | 0.39 | 0.50 |
| 2 | 99.94± | 98.97± | 99.69± | 98.75± | 98.23± | 99.72± | 98.82± | 99.41± | 98.83± | 97.28± |
|  | 0.27 | 1.85 | 1.32 | 1.01 | 1.31 | 0.34 | 2.72 | 1.87 | 2.06 | 0.48 |
| 3 | 97.72± | 97.93± | 98.76± | 98.79± | 98.71± | 98.54± | 99.12± | 98.00± | 97.21± | 98.34± |
|  | 1.05 | 1.31 | 2.27 | 1.97 | 2.82 | 0.69 | 0.67 | 1.47 | 0.51 | 1.43 |
| 4 | 98.68± | 98.62± | 97.76± | 95.24± | 95.82± | 99.95± | 99.74± | 98.62± | 98.72± | 97.73± |
|  | 0.36 | 0.47 | 1.33 | 1.38 | 1.10 | 0.17 | 0.37 | 0.72 | 0.66 | 0.40 |
| 5 | 100.13 | 98.85± | 98.34± | 98.61± | 97.96± | 99.70± | 99.47± | 99.02± | 96.69± | 96.29± |
|  | ±1.49 | 1.76 | 1.30 | 0.51 | 0.94 | 1.46 | 2.43 | 2.39 | 1.19 | 0.77 |
| 6 | 100.25 | 99.91± | 99.28± | 99.10± | 98.64± | 99.01± | 97.31± | 97.83± | 97.02± | 96.89± |
|  | ±1.13 | 1.76 | 1.12 | 1.22 | 0.81 | 0.90 | 1.14 | 1.96 | 0.80 | 1.53 |

## Discussion

In this study, the physicochemical compatibility and emulsion stability of urapidil injection and propofol emulsion were systematically evaluated under conditions for simulated Y-site co-administration. The results demonstrated excellent chemical compatibility between the two drugs for over 12 h. However, physical compatibility, particularly lipid emulsion stability, was robustly influenced by the mixing ratio, diluent type, and contact time. These findings provide vital clinical insights into the safe co-administration of these medications in critically ill ICU-admitted patients.

In this study, the relative contents of the two drugs in all admixtures remained above 95% throughout the 12-h observation period, and new impurity peaks were not detected. These results indicate chemical compatibility between urapidil and propofol, and contrast with those reported by Gersonde et al. [18], who observed ester bond hydrolysis and chemical degradation in propofol-remifentanil mixtures. Remifentanil contains an ester moiety susceptible to pH-dependent hydrolysis under mildly alkaline conditions, inactivating the drug. In contrast, urapidil, a phenylpiperazine derivative, lacks such labile functional groups and maintains chemical stability within the slightly acidic to near-neutral pH range (pH 6.11–6.74) of the admixtures [19,20].

To accurately detect potential chemical degradation, the HPLC conditions were systematically optimized. Considering the marked difference in acidity/basicity between urapidil (pKa ≈ 5.6) and propofol (pKa ≈ 11), the mobile phase pH was adjusted to 3.0. This condition enhanced urapidil ionization and peak shape, while suppressing propofol ionization, enabling satisfactory separation and detection of both drugs by the reversed-phase chromatographic system [20,21]. Furthermore, given the substantial difference in the polarities of the two analytes, a gradient elution program was employed, and the column temperature was maintained at 38 °C to improve mass transfer efficiency. Such an approach achieved baseline separation of the parent drugs and related impurities. The method established provides a useful reference for the quantitative analysis of similar multicomponent intravenous admixtures in future studies.

Despite satisfactory chemical stability, the physical stability assessment revealed potential safety concerns. The most critical finding was that, with an enhanced proportion of urapidil and 0.9% NS as the diluent, PFAT_5_ increased markedly after 2‒8 h and exceeded the pharmacopoeial safety limit of 0.05%. This phenomenon can be explained by the classical Derjaguin-Landau-Verwey-Overbeek theory [22]. The stability of propofol lipid emulsions largely depends on the electrostatic repulsion between oil droplets. As urapidil is administered as a hydrochloride salt, increasing its proportion within the NS-diluted solution introduced additional electrolytes—Na⁺ and Cl⁻—into the system. These cations compressed the electrical double layer at the droplet surface, significantly reducing the absolute zeta potential. The weakened electrostatic repulsion promoted droplet coalescence driven by van der Waals forces, forming large droplets (> 5 µm) [23]. The significant variations in zeta potential among groups (P < 0.001) provided direct experimental support for this mechanism.

Notably, in all samples with PFAT_5_ exceeding the safety limit, MDD stabilized at 169.1–182.7 nm across the study period, a finding consistent with an earlier report [24], suggesting that the microscopic aggregation of large lipid droplets may precede visible emulsion disruption or phase separation. These results further demonstrate that PFAT_5_ was more sensitive than MDD and served as an earlier indicator of emulsion destabilization, highlighting that MDD alone was insufficient for comprehensive assessment of lipid emulsion physical safety [25].

Another critical finding was the significant protective influence of 5% GS against emulsion destabilization induced by a high urapidil proportion. At the 1:10 mixing ratio, the NS group exceeded the PFAT_5_ safety limit, whereas the GS group stabilized over the 12-h study period. This effect may be attributed to the lack of electrolytes in GS, which prevents electrical double-layer compression. Additionally, the increased viscosity of the glucose-containing medium may reduce droplet collision and coalescence through steric hindrance.

These findings provide practical guidance for clinical use: when propofol and urapidil were co-administered via a Y-site infusion line, a 10:1 mixing ratio is recommended. If a greater proportion of urapidil is required, 5% GS has to be selected as the diluent. When NS must be used with an enhanced proportion of urapidil, the infusion duration should be < 2 h to reduce the potential risk of microembolism.

Several limitations of this study should be acknowledged. First, a static mixing model was used to simulate Y-site infusion, which may not fully reproduce the dynamic and transient mixing conditions prevalent at the clinical Y-site junction. In practice, the two drugs are rapidly mixed and infused into the bloodstream, with contact time typically limited to minutes. In contrast, the 12-h static model represents a worst-case scenario and may underestimate the risk compared with real-world clinical conditions. However, this conservative design helped identify potential risks associated with prolonged contact and provided a safety reference for situations such as infusion interruptions or rate fluctuations. Second, experiments were conducted at a constant room temperature (25 ± 2 °C) rather than body temperature. Elevated temperature may influence emulsion viscosity, droplet collision frequency, and drug degradation kinetics, thereby influencing compatibility and stability. Therefore, extrapolation to physiological conditions must be undertaken with caution. Finally, although representative mixing ratios were evaluated, the complexity of clinical infusion dynamics warrants further validation employing dynamic infusion models and in vivo pharmacokinetic studies.

## Conclusions

Under simulated Y-site administration conditions, urapidil and propofol demonstrated good chemical compatibility over 12 h, with no significant degradation. However, physical compatibility was affected by the mixing ratio, diluent type, and storage time. A propofol-dominant ratio (10:1, v/v) or the use of 5% GS maintained satisfactory emulsion stability. In contrast, increasing the urapidil proportion (1:1 or 1:10, v/v) with 0.9% NS as the diluent caused electrolyte-mediated reduction in zeta potential, causing PFAT_5_ to exceed the safety limit after 2‒8 h and indicating a potential risk of large-droplet embolism. This study provides new evidence regarding the compatibility and stability of urapidil-propofol admixtures. For clinical co-administration, a 10:1 mixing ratio or GS as a diluent is recommended. If a high proportion of NS-based urapidil solution is required, the infusion duration should be within 2 h.

## Supporting information

### S1 Fig. High-performance liquid chromatogram

(DOCX)

### S2 Raw data

(XLSX)

### Author contributions

**Conceptualization:** Yantao Yue, Liang Zhao.

**Data curation:** Yantao Yue, Shijie Feng.

**Methodology:** Liguo Wang, Yuanpei Li, Jingduo Zhao, Lijie Lou.

**Supervision:** Liang Zhao.

**Investigation:** Liguo Wang, Yuanpei Li.

**Writing—original draft:** Yantao Yue.

**writing—review & editing:** Liang Zhao, Yantao Yue.

### Funding

This research was funded by the Special Research Project of the National Health Commission Capacity Building and Continuing Education Center, grant number GWJJZX20251007028, and the Henan Provincial Medical Science and Technology Research Program, grant number LHGJ20240868, China.

## References

1. Custodio JEY, Navarro JEV, Malilay ORM. Conventional craniotomy and neuroendoscopic surgery for patients with hypertensive intracerebral hemorrhage: A meta-analysis and systematic review. Asian J Neurosurg. 2025;20: 478–484. 10.1055/s-0045-1809166 PMID: 40852060

2. Chinese Society of Neurology; Chinese College of Emergency Physicans; Chinese Stroke Society; Stroke Screening and Prevention Project Committee, National Health Commission of the People’s Republic of China. Chinese multidisciplinary guidelines for guideline for management of hypertensive intracerebral hemorrhage. Chin J Crit Care. 2020;40: 689–702.

3. Beijing Hypertension Association; Chinese Stroke Association Hypertension Prevention and Management Branch. Chinese Expert Consensus on Hypertension Management in Stroke Patients. Chin J Stroke. 2024;19: 672–698.

4. Cardiac Intensive Care Committee of China International Exchange and Promotive Association for Medical and Health Care; County Emergency and First Aid Group of the Emergency Medicine Branch of Chinese Medical Association; Primary Emergency Care Branch of the Emergency Medicine Branch of China Medicine Education Association; Multidisciplinary Expert Consensus Group on Clinical Application of Urapidil Hydrochloride Injection. Multidisciplinary expert consensus on the clinical application of urapidil hydrochloride injection. J Emerg Med. 2023;32: 305–313.

5. Devlin JW, Skrobik Y, Gélinas C, Needham DM, Slooter AJ, Pandharipande PP, et al. Clinical practice guidelines for the prevention and management of pain, agitation/sedation, delirium, immobility, and sleep disruption in adult patients in the ICU. Crit Care Med. 2018;46: e825–e873. 10.1097/ccm.0000000000003299 PMID: 30113379

6. Wu JB, Wang WH, Zhang Y, Hu LS, Luo F, Lin JM, et al. Significances of propofol in blood pressure control and prevention of re-hemorrhage after endoscopic surgery for hypertensive intracerebral hemorrhage. J Chin Pract Diagn Ther. 2021;35: 416–419.

7. Dong R, Li J, Chen Q, Zhang J, Zhang Z, Cao Y, et al. Variations in the clinical practice of blood pressure and sedo-analgesia management for intracerebral hemorrhage: A case-based survey of Chinese healthcare workers. BMC Neurol. 2025;25: 279–289. 10.1186/s12883-025-04301-6 PMID: 40618032

8. Koller AK, Krebs S, Dörje F. Medication safety in intravenous therapy: A compatibility study of clonidine with drugs frequently used in intensive care. Pharmaceutics. 2020;13: 21. 10.3390/pharmaceutics13010021 PMID: 33374362

9. Négrier L, Martin Mena A, Lebuffe G, Odou P, Genay S, Décaudin B. Strategies to prevent drug incompatibility during simultaneous multi-drug infusion in intensive care units: A literature review. Eur J Clin Pharmacol. 2021;77: 1309–1321. 10.1007/s00228-021-03112-1 PMID: 33768303

10. Keum N, Yoo J, Hur S, Shin S-Y, Dykes PC, Kang M-J, et al. The potential for drug incompatibility and its drivers—A hospital wide retrospective descriptive study. Int J Med Inform. 2024;191: 105584. 10.1016/j.ijmedinf.2024.105584 PMID: 39133962

11. Husna M, Munawiroh SZ, Puji Ekawati R, Hanifah S. Systematic review of the stability and compatibility of propofol injection. Anaesthesiol Intensive Ther. 2021;53: 79–88. 10.5114/ait.2021.103542 PMID: 33586420

12. Henkel E, Vella R, Behan K, Austin D, Kruger P, Fenning A. The effect of concentration, reconstitution solution and pH on the stability of a remifentanil hydrochloride and propofol admixture for simultaneous co-infusion. BMC Anesthesiol. 2020;20: 283–294. 10.1186/s12871-020-01194-5 PMID: 33183219

13. Klang MG. PFAT5 and the evolution of lipid admixture stability. J Parenter Enteral Nutr. 2015;39: 67S–71S. 10.1177/0148607115595976 PMID: 26177666

14. Gao SS, Fan J, Jia XL, Li BB, Li XC, Li ZL, et al. PFAT5 stability assessment of Lipovenoes MCT in total nutrient admixtures. Ann Palliat Med. 2021;10: 12244–12250. 10.21037/apm-21-3353 PMID: 35016476

15. Chinese Pharmacopoeia Commission. Chinese Pharmacopoeia (the Fourth Volume). Med Sci Technol Press China. 2020: 125–126.

16. Pharmacy Intravenous Admixture Services Professional Committee of Anhui Pharmacists Association; the First Affiliated Hospital of University of Science and Technology of China ( Anhui Provincial Hospital); the First Affiliated Hospital of Anhui University of Chinese Medicine. Expert consensus on physical stability and compatibility inspection indicators for ready-to-administer infusion solutions. Her Med. 2024;43: 149–155.

17. The United States pharmacopeia and national formulary. 38th ed. Rockville: United States Pharmacopeial Convention; 2015.

18. Gersonde F, Eisend S, Haake N, Kunze T. Physicochemical compatibility and emulsion stability of propofol with commonly used analgesics and sedatives in an intensive care unit. Eur J Hosp Pharm. 2017;24: 293–303. 10.1136/ejhpharm-2016-001038 PMID: 31156960

19. Zhao XY, Zhang LT, Yu SQ, Fan LJ, An J, Dong ZJ. Compatibility and stability study of butorphanol, nicardipine, urapidil, and tirofiban during multidrug simultaneous infusion. Drug Des Devel Ther. 2026; 20:578559 10.2147/dddt.s578559 PMID: 41804455

20. Zeng JT. Determination of propofol content and related substances in propofol injection by UPLC. Pharm. Clin Res. 2016;24: 131–133.

21. Roche M, Rousseleau D, Danel C, Henry H, Lebuffe G, Odou P, et al. Combination of a propofol emulsion with alpha-2 adrenergic receptor agonists used for multimodal analgesia or sedation in intensive care units: A physicochemical stability study. Eur J Hosp Pharm. 2025;32: 461–467. 10.1136/ejhpharm-2023-004027 PMID: 38290833

22. Zhang WQ, Jiang JZ, Cui ZG, Binks BP. Influence of positively charged nanoparticles on the stability of oil-in-water emulsions stabilized by a cationic surfactant at extremely low concentration. J Colloid Interface Sci. 2025;687: 449–460. 10.1016/j.jcis.2025.02.059 PMID: 39970585

23. Stawny M, Gostyńska A, Nadolna M, Jelińska A. Safe practice of Y-site drug administration: The case of colistin and parenteral nutrition. Pharmaceutics. 2020;12: 292. 10.3390/pharmaceutics12030292 PMID: 32213881

24. Nilsson N, Nezvalova-Henriksen K, Tho I. Emulsion stability of different intravenous propofol formulations in simulated co-administration with remifentanil hydrochloride. Pharm Technol Hosp Pharm. 2019;4: 77–87. 10.1515/pthp-2019-0014

25. Franco CP, Iglesias MU, Marchal AC, Polo AV, Muñoz PG, Berlana, D, et al. Drug compatibility with parenteral nutrition - Recommendations for evaluation and safe coadministration. Nutr Hosp. 2026; 43: 51–59. https://sci-hub.st/10.20960/nh.06653 PMID: 42023848

